# Distinct roles of microglial and neuronal P2X4 in persistent neuropathic pain

**DOI:** 10.64898/2026.11.19.767853

**Authors:** Damien Gilabert, Benjamin Sueur, Paola Nobili, Nathalie Linck, Pascal Fossat, Eric Boué-Grabot, Lauriane Ulmann

## Abstract

Neuropathic pain is a chronic, debilitating condition characterized by persistent sensory hypersensitivity and frequently associated with anxiety- and depression. Although microglial P2X4 receptors are well-established mediators of early neuropathic pain, their cell-specific contributions to pain maintenance and associated affective disturbances remains poorly understood. We investigated these contributions in male and female mice 30 and 60 days after spared nerve injury (SNI) using conditional P2X4 deletion in microglia (Cx3cr1CreERT2) or neurons (Synapsin-1-Cre), behavioral assessments, and *in vivo* recordings of spinal dorsal horn wide dynamic range neurons. Microglial P2X4 deletion before SNI completely prevented mechanical allodynia whereas deletion initiated 40 days after injury fully reversed established allodynia in both sexes. Notably, P2X4 immunoreactivity was undetectable in spinal microglia at the late time points examined, suggesting that the sustained contribution of microglial P2X4 may involve mechanisms outside the spinal microglial compartment. Neuronal P2X4 deletion partially attenuated mechanical hypersensitivity and reduced dorsal horn neurons hyperexcitability at day 60 after SNI, supporting a modulatory role in persistent spinal sensitization. Global P2X4 deletion prevented anxiety- and depressive-like behaviors that emerged by day 30 in males and day 60 in females. Whether this protection reflects a direct contribution of P2X4 to affective disturbances or an indirect consequence of reduced pain remains unresolved. Together, these findings identify distinct microglial and neuronal P2X4 contributions to established neuropathic pain and reveal sex-dependent differences in the onset of associated affective disturbances.

## Introduction

Neuropathic pain is a chronic and debilitating condition that remains inadequately managed by current therapies. Beyond persistent sensory hypersensitivity, chronic neuropathic pain is frequently associated with affective comorbidities, including anxiety and depression, which progressively emerge as pain persists and may contribute to its long-term burden and maintenance ^1^ ^2^. Importantly, sensory and affective manifestations do not necessarily follow the same temporal dynamics. Experimental studies have shown that anxiety- and depressive-like behaviors progressively develop after the establishment of sensory hypersensitivity ^2^, and sex differences have also been reported in the onset of pain-associated affective disturbances ^3^. These observations highlight the importance of considering both chronicity and sex when investigating the mechanisms underlying long-term neuropathic pain.

Among the molecular mechanisms involved in neuropathic pain, the purinergic receptor P2X4 has emerged as a key regulator of spinal excitability. Following peripheral nerve injury, P2X4 is strongly upregulated in spinal microglia, where ATP-dependent activation promotes BDNF release. Through neuronal TrkB signaling, BDNF downregulates KCC2, disrupting chloride homeostasis and weakening GABAergic and glycinergic inhibition. This disinhibition increases spinal dorsal horn excitability, contributing to mechanical hypersensitivity and tactile allodynia ^4^ ^5^ ^6^ ^7^. This P2X4-dependent microglia-to-neuron signaling pathway has been extensively characterized during the early phase of neuropathic pain, particularly within the first two weeks following nerve injury ^7^ ^8^.

However, whether P2X4 contributes to the maintenance of established neuropathic pain and which cellular populations mediate this contribution, remain unclear. Although sustained P2X4 upregulation has been reported for several weeks after central nervous system injury ^9^ and microglial P2X4 signaling has been implicated in chronic pain mechanisms ^10^, these findings do not establish its role at late stages following peripheral nerve injury. Moreover, while research has largely focused on microglial P2X4, the receptor is also expressed in sensory and spinal neurons, where it can modulate excitability and synaptic transmission ^11^ ^12^ ^13^. The respective contributions of microglial and neuronal P2X4 to persistent neuropathic pain and associated anxiety- and depressive-like behaviors remain be determined.

Here, we examined the respective contributions of microglial and neuronal P2X4 to established neuropathic pain in male and female mice at 30 and 60 days after spared nerve injury (SNI). These time points capture distinct stages of disease progression: at day 30 (D30), sensory hypersensitivity is firmly established in both sexes, whereas affective disturbances are detectable in males but not yet in females. By day 60, both sensory and affective alterations are established in both sexes. We combined conditional microglial or neuronal P2X4 deletion with behavioral assessments and *in vivo* recordings of spinal wide dynamic range neurons, and used global P2X4 knockout mice to examine associated affective disturbances. Our findings support a sustained contribution of microglial P2X4 despite undetectable spinal microglial expression at late stages, alongside a neuronal contribution to persistent spinal hyperexcitability. Global P2X4 deletion also prevented affective disturbances, which emerged at different times in males and females, although a direct role independent of pain remains unresolved.

## Methods

### Mice

Female and male mice from 6 to 10 weeks old were housed under a standard 12 h light/dark cycle with food and water available *ad libitum*. All the animals used were in C57BL/6J background. Mice carrying a null mutation for P2X4 gene (P2X4KO) and internalization-defective P2X4mCherryIN mice (P2X4KI) have been previously described^14^ ^15^ ^16^. To delete P2X4 expression in defined cell population, we used the floxed P2X4 KO lox/lox receptor obtained from Rieko Muramatsu’s Lab ^17^. To generate neuronal and microglial cell-specific P2X4 KO mice, with deletion of P2X4 selectively in microglia or neurons, we first generated double transgenic mice, by breeding P2X4^flox/flox^ with promoter specific-Cre mice expressing the Cre-recombinase under the control of neuronal Synapsin-1 (SYN-Cre) ^18^ or microglia Cx3cr1^CreERT^^2^ promoters ^19^.

The obtained heterozygote lines Cx3cr1CreERT2^+/wt^ : P2X4KO^+/flox^, or SYN-Cre^+/wt^ : P2X4KO^+/flox^ was crossed with P2X4^flox/flox^ to obtain homozygote Cx3cr1CreERT2^+/wt^ : P2X4KO^flox/flox^, or SYN-Cre^+/wt^ : P2X4KO^flox/flox^ ^19^ ^17^. All the mice were bred either in the SPF animal facility of the Institute for Functional Genomic (IGF, Montpellier, France; Agreement from the Ministry of Agriculture N° D34-172-13) or in the Pole In Vivo-Expe and EOPS production facilities of the Center Broca-Nouvelle Aquitaine of Bordeaux (Agreement from the Ministry of Agriculture are respectively N° A33-063-940 and N° A33-063-941).

### Spared nerve injury (SNI)

Mice were randomly assigned to SNI or sham group. Animals were anesthetized under 3% isoflurane/oxygen, the left leg sciatic nerve were exposed and the peroneal and tibial branches were ligated and transected. Skin was closed with silk sutures. Identical procedure was performed in sham operated mice except the nerve ligation and transection. The mice were isolated until they had fully recovered from the surgery before being reintroduced with their littermate^17^ ^20^.

### Tamoxifen

To induce recombination in Cx3cr1^CreERT2^ mice line, tamoxifen (T5648, Sigma) was dissolved at 10 mg/ml in a warm solution containing corn oil (C8267, Sigma) and 2.5% of absolute ethanol. Animals were injected with 1 mg of tamoxifen intraperitoneally one time a day during 3 days. Surgery was performed 21 days after the last injection allowing for peripheral CX3CR1+ cells replenishment while microglia remained depleted ^21^ ^22^.

### Tissue preparation

Mice were i.p euthanized with Euthasol Vet® (300 mg/kg) and perfused with PBS. Spinal cords were collected and fixed overnight in 4% paraformaldehyde. The contralateral side of the lesion was recognized by a scalpel’s notch. The lumbar segments were cut with a vibratome (VT1000S, Leica) into 40 μm sections and stored in PBS +0.1% sodium azide at 4°C.

### Immunohistochemistry

The sections were washed with PBS. Non-specific sites were saturated with 10% donkey serum (S30, Millipore) diluted in PBS +0.3% Triton X-100 (Sigma, X100) for 1h at room temperature. Primary antibodies (rabbit anti-Iba1, 1:2000, AB178846, Abcam, rat anti-mouse P2X4, a gift from Pr. Koch-Nolte^23^), were incubated overnight at 4°C in DPBS +0.3% Triton X-100. After three washes in PBS, the sections were incubated for 2h at room temperature with the corresponding secondary antibodies diluted in PBS +0.3% Triton X-100. The sections were rinsed with PBS 3 × 10 min and then mounted between slide and coverslip with mounting medium (Dako, S3023). L4 to L6 sections were observed using a fluorescence microscope (Axioimager 2, Zeiss) with an apotome grid module (Apotome 3, Zeiss) equipped with a digital camera (cool-snap HQ, PhotoMetrics) or confocal microscope (Leica SP8-UV).

### Mechanical hyperalgesia test

To test mechanical hyperalgesia, mice were habituated in Plexiglas compartments (5 cm × 10 cm x 14 cm) on a perforated metal floor for at least 45 min before the test. The hind paws withdrawal thresholds were measured on the different mouse lines in two distinct laboratories (Montpellier and Bordeaux) using electronic and manual Von Frey (Anesthesiometer, IITC Life Science, USA). For electronic Von Frey, calibrated tips were applied at the lateral plantar surface of the hind paw and response was recorded. Three measures per hind paw were recorded and averaged. For manual Von Frey (Ugo Basile, Italy), we calibrated Von Frey filament (in grams) on the plantar surface of the hind paws of the mice. Three to five measurements were made for each paw, with an interval of 30 s between each. The smallest grams of the filament at which the mouse withdrew its paw was taken as the mechanical pain threshold ^24^.

### Weight distribution changes test

Change in the hind paw weight distribution were assessed using the static weight-bearing test (WB), an incapacitance apparatus (BIO-SWB-TOUCH-M, Bioseb, France). Mice were placed in a Plexiglas box with an inclined plane, which permit to maintain animals in position. The weight applied by hind paws is measured during 10s. A mean of the weight over 10 s is recorded. Results were presented as ratio ipsilateral over contralateral to avoid influence of the animal body weight on the measurement.

### *In vivo* extracellular recordings

Mice were anesthetized with isoflurane inhalation and placed on a stereotaxic frame (Unimécanique, Asnières, France). Mice were first induced under 3% isoflurane/air before maintaining anesthesia during the experiment under 1.2% isoflurane/air. For the whole experiment, the air flow rate was set to 0.7 L/min. A laminectomy was performed on lumbar vertebrae L1–L3 and segments L4–L5 of the spinal cord were exposed. Extracellular recordings of wide dynamic range (WDR) dorsal horn neurons were made with borosilicate glass capillaries (2 M*Ω*, filled with NaCl 684 mM) (Harvard Apparatus, Cambridge, MA, USA). The criterion for the selection of a neuron was the presence of an A-fiber-evoked response (0-80 ms) followed by a C fiber-evoked response (80-300 ms) to electrical stimulation of the ipsilateral paw with subcutaneous bipolar electrodes connected to a stimulator and placed in the center of the recorded neuron receptive field. Threshold for C-fiber evoked response was evaluated by progressive increase in electrical stimulation intensity until the appearance of a C-fiber component. Intensity response curved were measured with incremental stimulations from 1mA to 5 mA. The number of C-spikes was directly measured and plotted against stimulation intensity.

All measured are expressed as mean ± SEM. WDR threshold in each experimental group was compared using two-way ANOVA followed with Tukey’s post hoc analysis p value < 0.05 was considered significant.

### Quantification and statistical analysis

Data are presented as mean ± SEM. Statistical analyses were performed using GraphPad Prism. Groups were compared using two-way ANOVA followed with Tukey’s post hoc analysis p value < 0.05 was considered significant ^∗^p < 0.05, ∗∗p < 0.01, ∗∗∗p < 0.001, ∗∗∗∗p < 0.0001, ns p > 0.05.

## Results

### Microglial P2X4 upregulation in the spinal cord is transient after SNI

To determine whether spinal microglial P2X4 expression persists during established neuropathic pain, we performed double immunostaining for P2X4 and the microglial marker Iba1 in the ipsilateral dorsal horn of the lumbar spinal cord at 15, 30 and 60 days after SNI. P2X4 immunoreactivity was markedly increased in Iba1-positive microglia at D15 after SNI but returned to baseline levels by D30, despite persistent morphological signs of microglial reactivity (**Fig. 1A**). These findings indicate that SNI induces a transient upregulation of P2X4 in spinal microglia.

**Figure 1.**
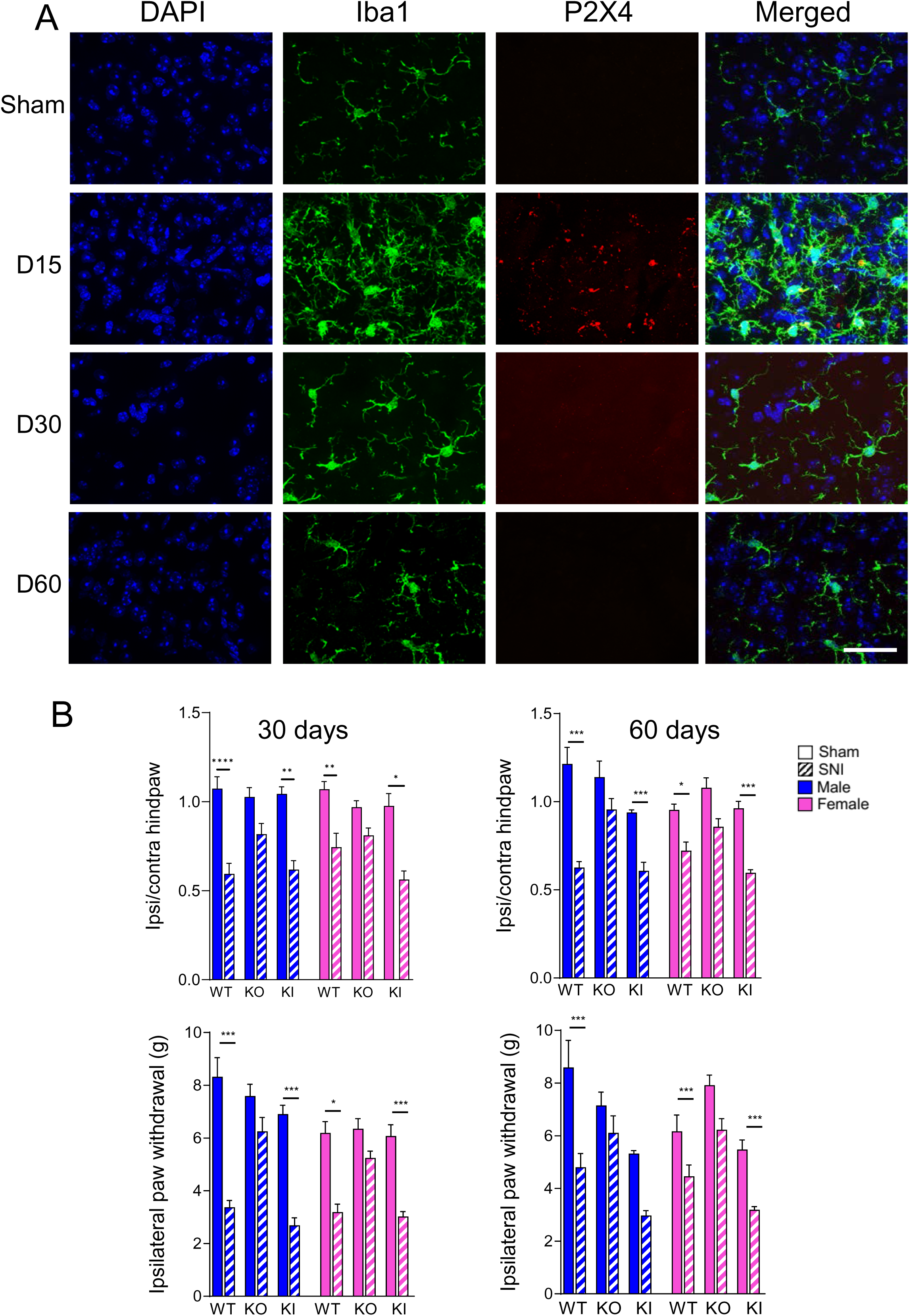
Transient spinal microglial P2X4 upregulation and persistent mechanical hypersensitivity after SNI. **(A)** Representative confocal images of the ipsilateral dorsal horn of the L4-L6 lumbar spinal cord co-immunostained for P2X4 (red), the microglial marker Iba1 (green) and DAPI (blue), in sham-operated mice and at 15, 30 and 60 days after spared nerve injury. Scale bar = 50 µm. **(B)** Mechanical sensitivity at D30 and D60 after SNI in wild-type (WT), internalization-defective P2X4mCherryIN knock-in (KI) and constitutive P2X4 knock-out (KO) mice of both sexes. Left panel, weight-bearing (WB) expressed as the ipsilateral/contralateral hind paw weight ratio. Right panel, paw withdrawal threshold (g) measured by von Frey. WT and KI SNI mice show a significant deficit relative to their respective sham controls at D30 that is unchanged at D60 in both measures, whereas P2X4KO SNI mice are indistinguishable from sham at both timepoints. Group means ± SEM are reported in Table 1. Data are represented as mean ± SEM; *n* = 6-17 mice per group. Three-way ANOVA followed by FDR-corrected post hoc comparisons; ∗*p* < 0.05, ∗∗*p* < 0.01, ∗∗∗*p* < 0.001, ∗∗∗∗*p* < 0.0001.

**Table 1:** Pain-related behaviors in male and female WT, P2X4KO and KI mice 30 and 60 days after sham or SNI surgery. Static weight distribution was assessed with the weight-bearing test (WB; ipsilateral/contralateral hind paw weight ratio) and mechanical sensitivity with the dynamic von Frey test (VF; paw withdrawal threshold, g). Values are expressed as mean ± SEM. n = 6-17 mice per group. D30/D60, days post-surgery; SNI, spared nerve injury; WT, wild type; P2X4KO, P2X4 knockout; KI, P2X4 knock in mouse.

|  |  | **Males** |  |  |  |  |  | **Females** |  |  |  |  |  |
|---|---|---|---|---|---|---|---|---|---|---|---|---|---|
|  |  | **WT sham** | **WT SNI** | **P2X4KO sham** | **P2X4KO SNI** | **KI sham** | **KI SNI** | **WT sham** | **WT<br>SNI** | **P2X4KO sham** | **P2X4KO SNI** | **KI sham** | **KI SNI** |
| **WB** | D30 | 1.07 ± 0.07 | 0.59 ± 0.06 | 1.03 ± 0.05 | 0.82 ± 0.06 | 1.04 ± 0.04 | 0.62 ± 0.05 | 1.07 ± 0.04 | 0.74 ± 0.08 | 0.97 ± 0.03 | 0.81 ± 0.04 | 0.98 ± 0.07 | 0.56 ± 0.05 |
|  | D60 | 1.21 ± 0.09 | 0.63 ± 0.03 | 1.14 ± 0.09 | 0.96 ± 0.06 | 0.94 ± 0.01 | 0.61 ± 0.05 | 0.95 ± 0.03 | 0.72 ± 0.05 | 1.08 ± 0.06 | 0.86 ± 0.05 | 0.96 ± 0.04 | 0.60 ± 0.02 |
| **VF** | D30 | 8.32 ± 0.73 | 3.37 ± 0.26 | 7.59 ± 0.44 | 6.25 ± 0.53 | 6.91 ± 0.33 | 2.69 ± 0.29 | 6.20 ± 0.43 | 3.19 ± 0.30 | 6.35 ± 0.39 | 5.25 ± 0.26 | 6.07 ± 0.43 | 3.02 ± 0.19 |
|  | D60 | 8.60 ± 1.03 | 4.81 ± 0.52 | 7.15 ± 0.51 | 6.11 ± 0.64 | 5.33 ± 0.11 | 2.98 ± 0.18 | 6.17 ± 0.61 | 4.46 ± 0.42 | 7.92 ± 0.38 | 6.23 ± 0.42 | 5.48 ± 0.36 | 3.19 ± 0.12 |

To investigate whether increased P2X4 cell-surface availability exacerbates mechanical hypersensitivity at late stages after SNI, we examined wild-type (WT) and internalization-defective P2X4 knock-in (KI) mice, alongside global P2X4 knockout (KO) mice (**Fig. 1B**; **Table 1**). At D30, both WT and KI SNI mice exhibited a significantly reduced ipsilateral-to-contralateral hindlimb weight-bearing ratio compared with their respective sham controls, indicating reduced loading of the injured hindlimb. Dynamic von Frey testing revealed pronounced mechanical allodynia in both genotypes at D30, with significantly lower paw withdrawal thresholds than in their respective sham groups. This hypersensitivity persisted at D60. In contrast, withdrawal thresholds in P2X4KO SNI mice did not differ significantly from those of sham WT at either time point. Together, these findings support a contribution of P2X4 to persistent mechanical hypersensitivity, despite the transient nature of its detectable upregulation in spinal microglia.

### Anxiety- and depressive-like behaviors develop earlier in males than in females

Given the frequent association of chronic pain with anxiety and depression ^25^, we next assessed anxiety- and depressive-like behaviors in male and female mice at 30 and 60 days after SNI.

To control for potential locomotor confounds, locomotor activity was assessed using the Cyclotron test. No significant differences were detected between SNI and sham groups across genotypes, sexes or time points (**Fig. 2A** and **Table 2**) suggesting that the behavioral findings were unlikely to reflect changes in general locomotor activity.

**Figure 2.**
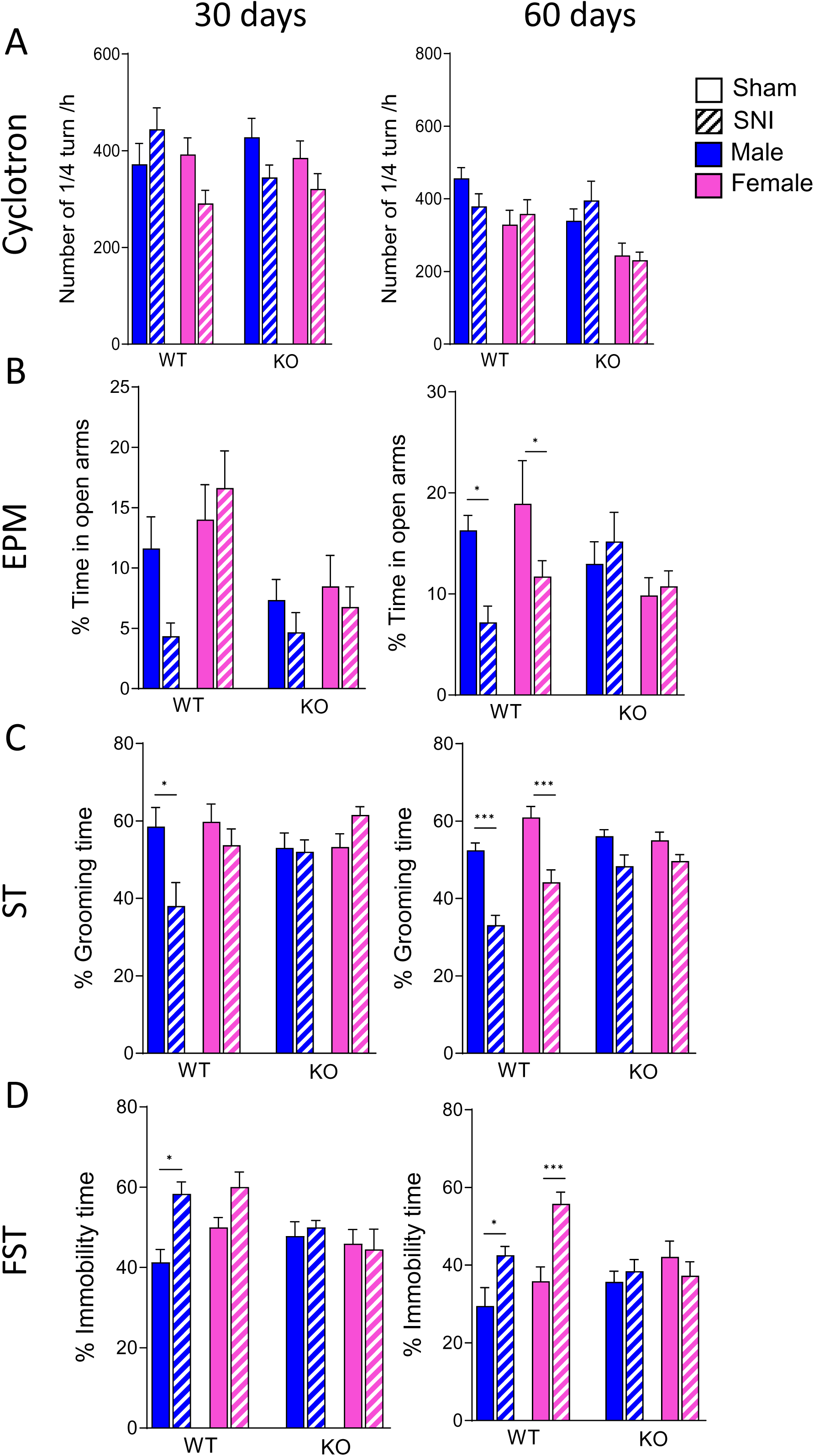
SNI-induced anxiety and depressive-like behaviors develop earlier in males than in females and are absent in P2X4KO mice. **(A)** Locomotor activity in the cyclotron test, expressed as the number of ¼ turn/h. No effect of surgery, genotype or sex is detected at either timepoint, indicating that the differences reported in **(B-D)** are not secondary to altered motor activity. **(B)** Anxiety-like behavior in the elevated plus maze (EPM), expressed as the percentage of time spent in the open arms. WT males show a significant reduction after SNI from D30 onwards, sustained at D60; WT females show no difference at D30 but a significant reduction at D60. P2X4KO mice of either sex are unaffected at both timepoints. **(C)** Self-care behavior in the splash test (ST), expressed as total grooming time (s). WT males show a significant reduction from D30, sustained at D60; WT females only at D60. P2X4KO mice of either sex are unaffected. **(D)** Behavioral despair in the forced swim test (FST), expressed as immobility time (s). WT males show a significant increase from D30, sustained at D60; WT females only at D60. P2X4KO mice of either sex are unaffected. Group mean ± SEM for all four tests is reported in Table 2. Data are represented as mean ± SEM; n = 7-17 mice per group. Three-way ANOVA followed by FDR-corrected post hoc comparisons; ∗p < 0.05, ∗∗p < 0.01, ∗∗∗p < 0.001, ∗∗∗∗p < 0.0001; ns, not significant.

**Table 2:** Anxio-depressive behavioral characterization of male and female WT and P2X4KO mice 30 and 60 days after sham or SNI surgery. Locomotor activity was assessed in the cyclotron (number of ¼ turn/h), anxiety-like behavior in the elevated plus maze (EPM; % time spent in open arms), and depressive-like behavior in the splash test (ST; grooming time, s) and the forced swim test (FST; immobility time, s). Values are expressed as mean ± SEM. n = 6-17 mice per group. D30/D60, days post-surgery; SNI, spared nerve injury; WT, wild type; P2X4KO, P2X4 knockout.

|  |  | Males |  |  |  | Females |  |  |  |
|---|---|---|---|---|---|---|---|---|---|
|  |  | **WT sham** | **WT SNI** | **P2X4KO sham** | **P2X4KO SNI** | **WT sham** | **WT SNI** | **P2X4KO sham** | **P2X4KO SNI** |
| **Cyclotron** | D30 | 372 ± 43 | 444 ± 44 | 427 ± 39 | 344 ± 26 | 392 ± 34 | 291 ± 27 | 385 ± 35 | 321 ± 32 |
|  | D60 | 457 ± 29 | 379 ± 35 | 339 ± 33 | 395 ± 53 | 329 ± 40 | 359 ± 39 | 244 ± 34 | 231 ± 22 |
| **EPM** | D30 | 11.6 ± 2.6 | 4.3 ± 1.1 | 7.3 ± 1.7 | 4.7 ± 1.6 | 14.0 ± 2.9 | 16.6 ± 3.1 | 8.5 ± 2.6 | 6.8 ± 1.7 |
|  | D60 | 16.3 ± 1.5 | 7.2 ± 1.6 | 13.0 ± 2.2 | 15.2 ± 2.9 | 18.9 ± 4.3 | 11.7 ± 1.5 | 9.9 ± 1.8 | 10.8 ± 1.5 |
| **ST** | D30 | 58.5 ± 4.9 | 38.0 ± 6.1 | 53.0 ± 3.9 | 52.0 ± 3.1 | 59.8 ± 4.6 | 53.7 ± 4.2 | 53.2 ± 3.4 | 61.5 ± 2.2 |
|  | D60 | 52.4 ± 1.9 | 33.1 ± 2.5 | 56.1 ± 1.7 | 48.4 ± 2.9 | 61.0 ± 2.8 | 44.2 ± 3.2 | 55.1 ± 2.1 | 49.7 ± 1.7 |
| **FST** | D30 | 41.3 ± 3.2 | 58.4 ± 2.9 | 47.8 ± 3.6 | 50.0 ± 1.7 | 50.0 ± 2.5 | 60.0 ± 3.7 | 45.9 ± 3.5 | 44.5 ± 5.0 |
|  | D60 | 29.5 ± 4.7 | 42.6 ± 2.2 | 35.7 ± 2.8 | 38.4 ± 3.0 | 35.9 ± 3.6 | 55.8 ± 3.0 | 42.1 ± 4.1 | 37.3 ± 3.6 |

In the elevated plus maze (EPM), WT males, exhibited a significant reduction in open-arm exploration at D30 after SNI, consistent with anxiety-like behavior which persisted at D60 **(Fig. 2B)**. In WT females, no significant difference was observed at D30. whereas open-arm exploration was significantly reduced at D60. In contrast, P2X4KO mice of both sexes showed no significant SNI-induced changes at either time point.

In the splash test, WT males displayed a significant reduction in grooming time at D30, which persisted at D60 **(Fig. 2C)**. WT females showed no significant change at D30 but exhibited significantly reduced grooming at D60. In contrast, P2X4KO mice of both sexes showed no significant SNI-induced reduction of motivational/self-care at either time point.

Finally, in the forced swim test, SNI significantly increased immobility time in WT males at D30, and this increase persisted at D60 **(Fig. 2D)**. WT females exhibited no significant change at D30 but showed a significant increased immobility at D60. No significant SNI-induced changes were detected in P2X4KO mice of either sex at either time point.

Together, these results indicate that SNI-induced anxiety- and depressive-like behaviors were detectable earlier in WT males (D30) than in WT females (D60), but both sexes show similar alterations by D60. As expected, these behavioral changes were absent in P2X4KO mice.

### Preventive and delayed conditional deletion of microglial P2X4 prevents and reverses persistent pain hypersensitivity in both sexes

To determine whether microglial P2X4 is required for the development of persistent neuropathic pain, we used Cx3cr1^CreERT2^ : P2X4KO^flox/flox^ mice, in which tamoxifen administration induces conditional P2X4 deletion in microglia. Tamoxifen was administered 21 days before SNI, and behavioral tests were performed at D30 and D60 after injury in male and female mice.

In vehicle-treated control mice, SNI induced significant mechanical allodynia that persisted at D60, with markedly reduced paw withdrawal thresholds compared with sham controls at both time points (**Fig. 3A**, **Table 3**). In contrast, tamoxifen-treated Cx3cr1^CreERT2^ : P2X4KO^flox/flox^ mice (lacking microglial P2X4) showed no significant mechanical allodynia at either time point, with withdrawal thresholds comparable to those of sham controls. This protective effect was observed in both males and females. Similarly, the weight-bearing test revealed a significant reduction in ipsilateral hindlimb loading in control SNI mice at D30 and D60 (**Fig. 3B**). Tamoxifen-treated Cx3cr1^CreERT2^ : P2X4^flox/flox^ mice showed no weight-bearing deficit at either time point. These findings support a requirement for microglial P2X4 in the development of mechanical persistent allodynia and pain-related weight-bearing deficits in both sexes. This preventive effect is consistent with our previous observation that microglial P2X4 deletion before SNI prevents early allodynia ^17^, and extends this finding by showing that protection persists to D60 in both sexes.

**Figure 3:**
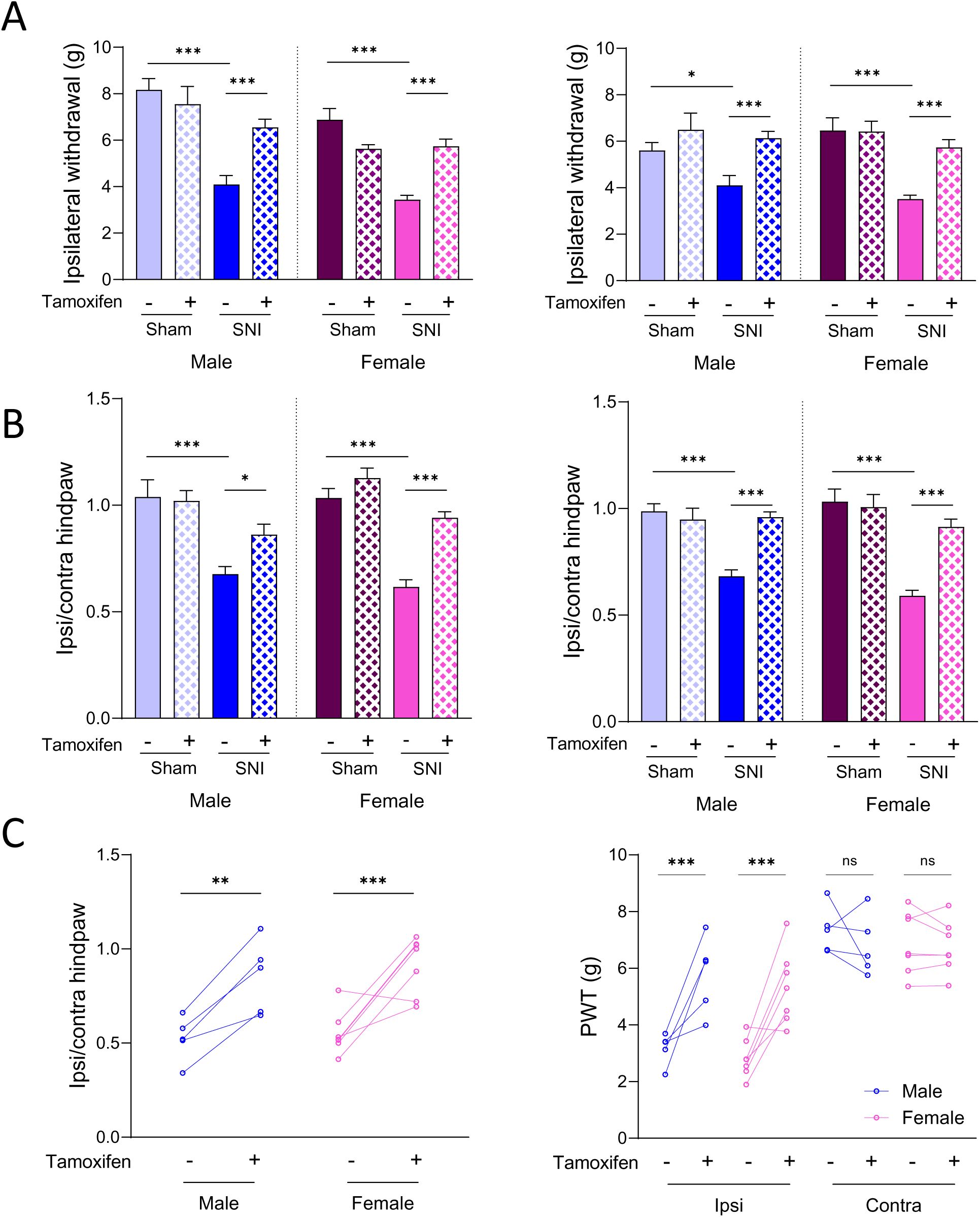
Preventive and delayed conditional P2X4 deletion attenuates persistent hypersensitivity in both sexes. Cx3cr1^CreERT2/+^ : P2X4^flox/flox^ mice received tamoxifen to delete P2X4 from the myeloid compartment, either before (A, B) or after (C) spared nerve injury (SNI). **(A, B)** Preventive deletion. Tamoxifen was administered 21 days before SNI and mice were tested at D30 (left panel) and D60 (right panel) post-injury. **(A)** Hind paw withdrawal threshold (g, von Frey). **(B)** Weight-bearing ratio (ipsilateral/contralateral). In vehicle-treated mice, SNI produced a significant deficit in both measures at D30 that persisted to D60. In tamoxifen-treated mice, SNI produced no significant deficit at either timepoint, in either sex. Group means ± SEM are reported in Table 3. **(C)** Delayed deletion. Tamoxifen was administered 40 days after SNI and mice were tested at D30 and D60. Left Panel, weight-bearing ratio before (Tamoxifen -, D30) and 20 days after tamoxifen injection (Tamoxifen +, D60); Right panel, hind paw withdrawal threshold (PWT, g, von Frey) before (Tamoxifen -, D30) and 20 days after tamoxifen injection (Tamoxifen +, D60). Tamoxifen-treated SNI mice showed a significant reversal of both mechanical allodynia and weight-bearing deficit. The effect was comparable in males and females. Data are represented as mean ± SEM; n = 5-20 mice per group. Two-way or Three-way ANOVA followed by FDR-corrected post hoc comparisons; ∗p < 0.05, ∗∗p < 0.01, ∗∗∗p < 0.001, ∗∗∗∗p < 0.0001.

**Table 3:** Effect of preventive deletion of microglial P2X4 on mechanical sensitivity and weight bearing in male and female mice 30 and 60 days after sham or SNI surgery. . *Cx3cr1*^CreERT2/+^ : P2X4KO^flox/flox^ mice received vehicle or tamoxifen 21 days before sham or SNI surgery. Mechanical sensitivity was assessed with the von Frey test (VF; ipsilateral hind paw withdrawal threshold, g) and static weight distribution with the weight-bearing test (WB; ipsilateral/contralateral hind paw weight ratio). Values are expressed as mean ± SEM; n = 5-20 mice per group. D30/D60, days post-surgery; SNI, spared nerve injury; Veh, vehicle; Tam, tamoxifen.

|  | **Males** |  | **Females** |  |
|---|---|---|---|---|
|  | **Before Tam (D30)** | **After Tam (D60)** | **Before Tam (D30)** | **After Tam (D60)** |
| **WB** | 0.52 ± 0.05 | 0.85 ± 0.09 | 0.55 ± 0.04 | 0.91 ± 0.06 |
| **VF ipsilateral** | 3.17 ± 0.25 | 5.76 ± 0.60 | 2.82 ± 0.26 | 5.34 ± 0.49 |
| **VF contralateral** | 7.35 ± 0.37 | 6.80 ± 0.48 | 6.88 ± 0.42 | 6.75 ± 0.35 |

To determine whether microglial P2X4 also contributes to the maintenance of established chronic pain, we administered tamoxifen at D40 after SNI and assessed behavioral responses at D60 (**Fig. 3C and Table 4**). In vehicle-treated control mice, mechanical allodynia and weight-bearing deficits persisted at D60. In contrast, tamoxifen-induced P2X4 deletion at D40 significantly reversed these alterations in both males and females with mechanical withdrawal thresholds and weight-bearing values approaching those of sham controls.

**Table 4:** Effect of delayed deletion of microglial P2X4 on weight bearing and mechanical sensitivity in male and female SNI mice. *Cx3cr1*^CreERT2/+^ : P2X4^flox/flox^ mice received tamoxifen 40 days after SNI. Static weight distribution was assessed with the weight-bearing test (WB; ipsilateral/contralateral hind paw weight ratio) and mechanical sensitivity of the ipsilateral and contralateral hind paws with the von Frey test (VF; paw withdrawal threshold, g), before (D30) and 20 days after tamoxifen injection (D60). Values are expressed as mean ± SEM; n = 5-7 mice per group. SNI, spared nerve injury; Tam, tamoxifen.

|  |  | **Males** |  |  |  | **Females** |  |  |  |
|---|---|---|---|---|---|---|---|---|---|
|  |  | **Sham Veh** | **Sham Tam** | **SNI Veh** | **SNI Tam** | **Sham Veh** | **Sham Tam** | **SNI Veh** | **SNI Tam** |
| **VF** | D30 | 8.17 ± 0.48 | 7.55 ± 0.76 | 4.10 ± 0.38 | 6.55 ± 0.35 | 6.88 ± 0.48 | 5.63 ± 0.17 | 3.43 ± 0.19 | 5.74 ± 0.30 |
|  | D60 | 5.60 ± 0.34 | 6.50 ± 0.72 | 4.10 ± 0.43 | 6.14 ± 0.29 | 6.46 ± 0.55 | 6.42 ± 0.44 | 3.51 ± 0.17 | 5.74 ± 0.33 |
| **WB** | D30 | 1.04 ± 0.08 | 1.02 ± 0.05 | 0.68 ± 0.04 | 0.86 ± 0.05 | 1.03 ± 0.04 | 1.13 ± 0.05 | 0.62 ± 0.03 | 0.94 ± 0.03 |
|  | D60 | 0.99 ± 0.04 | 0.95 ± 0.05 | 0.68 ± 0.03 | 0.96 ± 0.02 | 1.03 ± 0.06 | 1.01 ± 0.06 | 0.59 ± 0.03 | 0.91 ± 0.04 |

Together, these results indicate that microglial P2X4 contributes to both the development and maintenance of persistent neuropathic pain: deletion before SNI prevents long-term hypersensitivity, whereas deletion initiated after pain is established reverses hypersensitivity by D60 in both sexes.

### Neuronal P2X4 deletion attenuates persistent mechanical hypersensitivity and spinal neuronal hyperexcitability

Our immunohistochemical data showed that SNI-induced P2X4 upregulation in spinal microglia was no longer detectable at D30 or D60, despite persistent mechanical allodynia. To investigate whether neuronal P2X4 contribute to pain at these later stages, we used mice with conditional P2X4 deletion in neuronal populations (SynP2X4KO) generated by crossing P2X4KO^flox/flox^with synapsin-Cre mice (see methods).

We first assessed mechanical sensitivity using the von Frey test in sham and SNI mice. In sham males, WT and SynP2X4KO mice exhibited comparable mechanical withdrawal threshold throughout the 60-day observation period, with no significant change between baseline (before surgery) and D60 (**Fig. 4A**, upper panels, **Table 5**). Similar results where obtain in females (**Fig. 4A**, lower panels). In males, SNI significantly reduced mechanical withdrawal threshold in both genotypes. However, SynP2X4KO mice exhibited significantly less mechanical allodynia than WT mice at all post-SNI time points examined. Nevertheless, withdrawal thresholds remained significantly lower at D60 than at baseline in both genotypes (**Fig 4B**, upper panels). Similar effects were observed in females (**Fig. 4B**, lower panels). These findings indicate that neuronal P2X4 deletion attenuates, but does not prevent, persistent mechanical allodynia in either sex.

**Figure 4:**
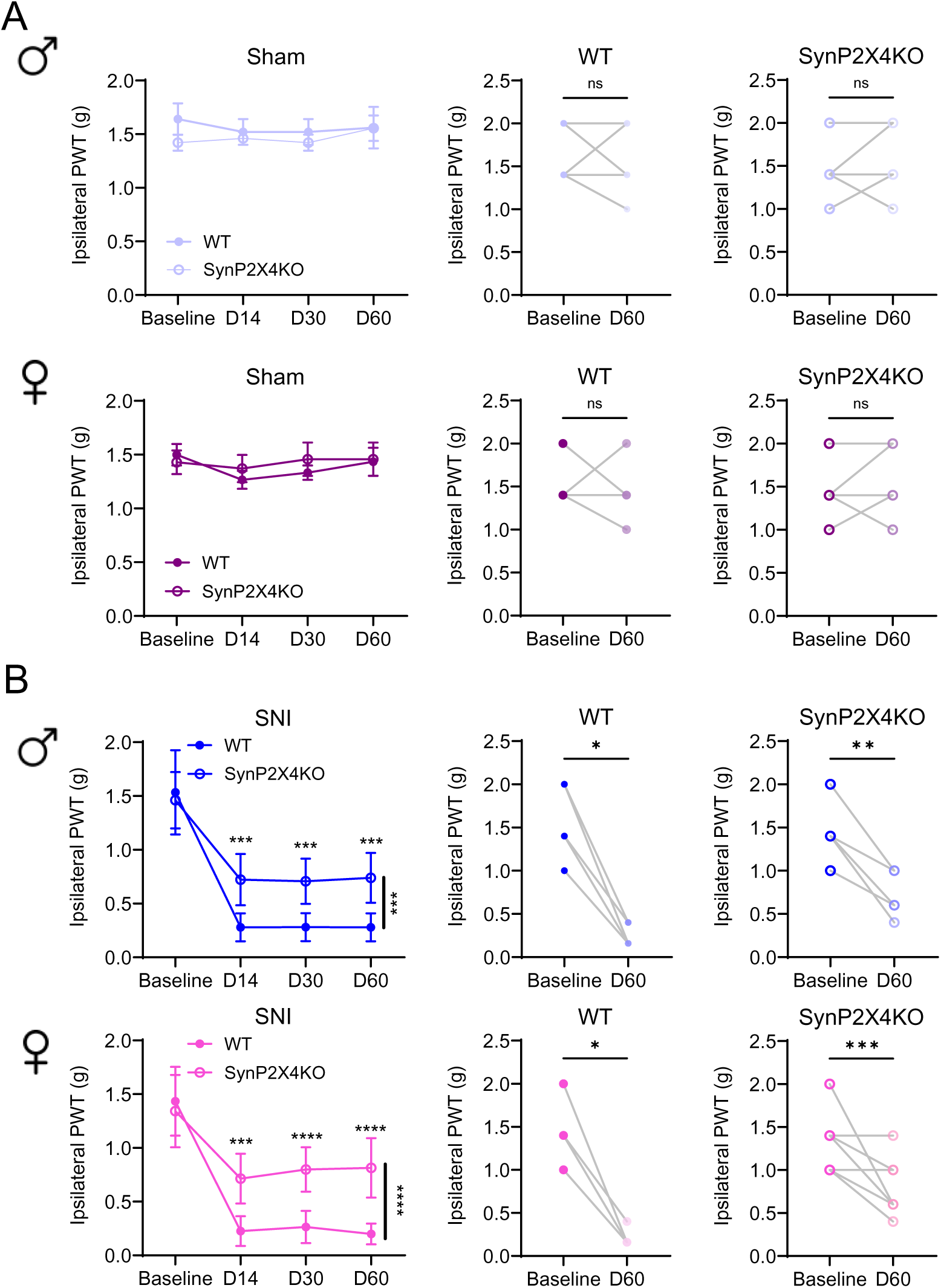
Neuronal P2X4 deletion partially attenuates SNI-induced mechanical allodynia in both sexes. **(A-B)** Manual Von Frey for the assessment of mechanical sensitivity of both sexes on WT (filled circle) and Neuronal P2X4KO (empty circle mouse lines on Sham (**A**; male: *bottom panels* in light blue; female: *top panels* in dark purple) and SNI (**B**; male: *bottom panels* in dark blue; female *top panels* in pink) conditions over time (assessment at 15, 30, and 60 days after surgery). A comparison between the baseline threshold and 60 days post-surgery for each animal has been performed. **(A)** In physiological condition the depletion of P2X4 in neurons in both males and females doesn’t change the mechanical sensitivity threshold due to an evoked nociceptive stimulus. **(B)** As expected, SNI induces a significant decrease in the mechanical paw withdrawal threshold in both male and female WT mice. Although also present, this observed decrease is less pronounced in neuronal P2X4KO mice. The neuronal P2X4KO attenuates the mechanical hypersensitivity induced by neuropathic pain suggesting a key role of neuronal P2X4 on neuropathic but not on physiological condition. (**A-B;** *left panels*) Data are represented as mean ± SEM, n=6-14 per condition, with mixed-effects analysis with Geisser-Greenhouse correction, followed by Šídák’s multiple comparisons test. (**A-B;** *central and right panels*) Data are represented as individual paired values (Baseline and 60 days post-surgery) of each animal using Wilcoxon test. **p<0.01, *** p <0.001, ****p<0.0001

**Table 5:** Mechanical sensitivity in male and female WT and Neuronal P2X4KO 15, 30 and 60 days after sham or SNI surgery. Mechanical sensitivity was assessed using a manual von Frey test (VF; paw withdrawal threshold, g). Values are expressed as mean ± SEM. n = 5-14 mice per group. D14/D30/D60, days post-surgery; SNI, spared nerve injury; WT, wild type; SynP2X4KO, neuronal P2X4 knockout mice.

| **VF behavior** | **Male** |  |  |  | **Female** |  |  |  |
|---|---|---|---|---|---|---|---|---|
|  | **Sham** |  | **SNI** |  | **Sham** |  | **SNI** |  |
|  | **WT** | **SynP2X4KO** | **WT** | **SynP2X4KO** | **WT** | **SynP2X4KO** | **WT** | **SynP2X4KO** |
| Baseline | 1.64 ± 0.14 | 1.42 ± 0.07 | 1.53 ± 0.16 | 1.46 ± 0.07 | 1.5 ± 0.1 | 1.42 ± 0.11 | 1.43 ± 0.13 | 1.34 ± 0.09 |
| D14 | 1.52 ± 0.12 | 1.46 ± 0.06 | 0.28 ± 0.05 | 0.72 ± 0.06 | 1.26 ± 0.08 | 1.37 ± 0.12 | 0.22 ± 0.05 | 0.71 ± 0.06 |
| D30 | 1.52 ± 0.12 | 1.42 ± 0.075 | 0.28 ± 0.05 | 0.70 ± 0.05 | 1.33 ± 0.06 | 1.45 ± 0.15 | 0.26 ± 0.06 | 0.8 ± 0.05 |
| D60 | 1.56 ± 0.19 | 1.44 ± 0.11 | 0.28 ± 0.05 | 0.74 ± 0.07 | 1.43 ± 0.13 | 1.45 ± 0.15 | 0.2 ± 0.04 | 0.81 ± 0.07 |

To investigate whether this attenuation was associated with reduced spinal neuronal excitability, we performed *in vivo* recordings of dorsal horn wide dynamic range (WDR) neurons at D60 in male and female WT and SynP2X4KO mice following sham surgery or SNI (**Fig. 5A**, **table 6**). We first measured the electrical stimulation threshold required to evoke a C-fiber-mediated response. In sham males, this threshold was slightly but significantly higher in SynP2X4KO mice than in WT mice and the difference between genotypes was more pronounced following SNI (**Fig. 5B**, upper panels). In females, no significant genotype difference was detected in sham mice, whereas SynP2X4KO SNI mice exhibited significantly higher thresholds than WT SNI mice (**Fig. 5B**, lower panels).

**Figure 5:**
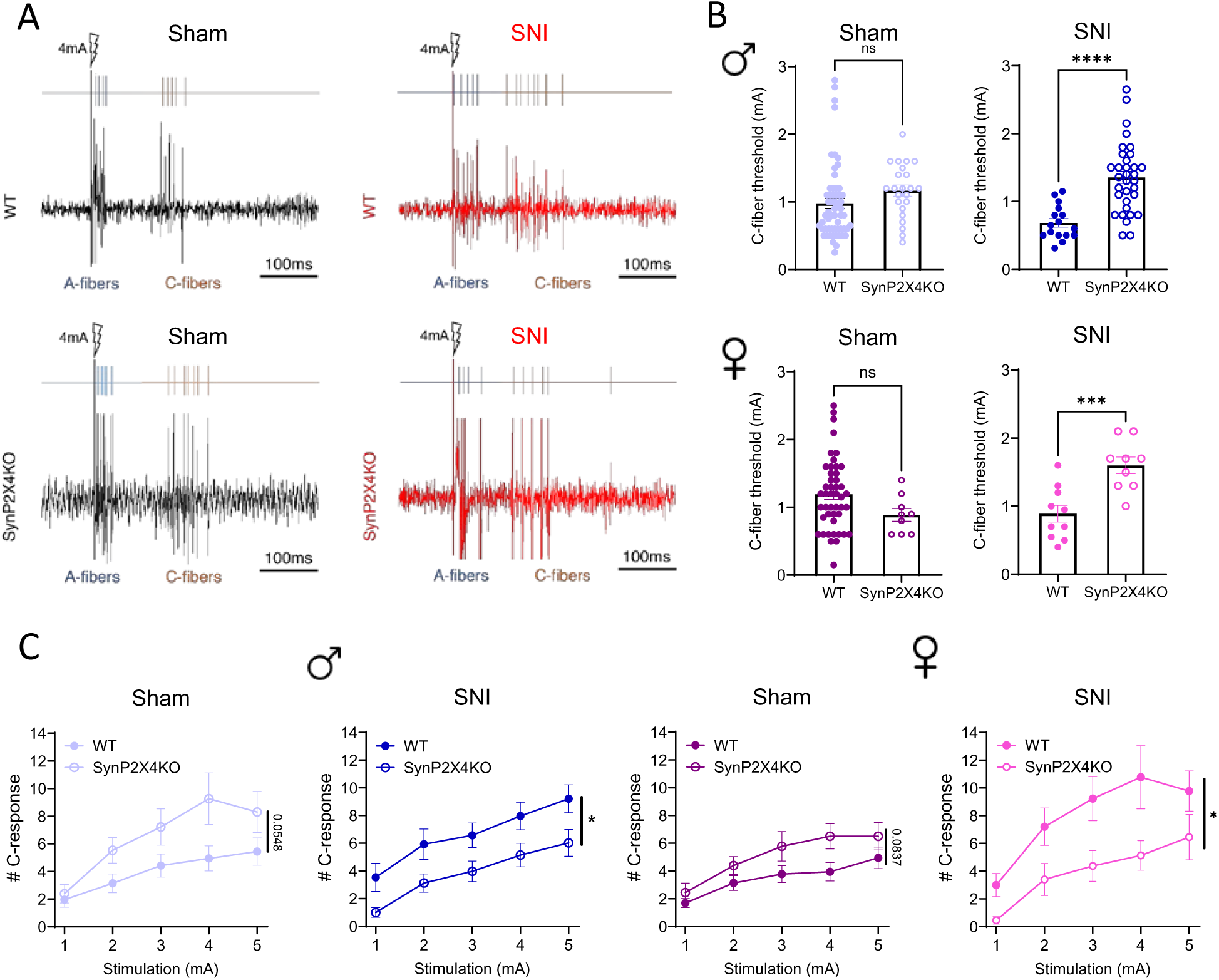
Neuronal P2X4 deletion reduces spinal WDR neuronal excitability after SNI. **(A-C)** *In vivo* single-unit recordings of WDR neurons recorded on the dorsal horn (L4-L5 segments) of the spinal cord in response to C-fiber stimulation triggered by electrical stimulation of the hindpaw of anesthetized mice. Data are represented as mean ± SEM n=9-58 per condition *p<0.05, *** p <0.001, ****p<0.0001. **(A)** Examples of action potentials recorded in Sham (*left panels*) or SNI (*right panels*) at 4mA stimulation on WT (black traces) and neuronal P2X4KO (red traces). Note that in SNI, a high-intensity stimulation of C-fibers induces a stronger WDR response in WT mice than in neuronal P2X4KO mice. **(B)** The C-fiber threshold is the minimum intensity of the peripheral receptive field required to elicit a response in WDR neurons, with a latency of 80-300 ms (*cf.* orange spikes on A panels). Compared to WT mice, under physiological conditions, the neuronal P2X4 knock-out slightly increases the C-fiber threshold in males but not in females, where the threshold tends to decrease. As previously described in the neuropathic condition, using the SNI procedure in WT animals, a significant decrease in C-fiber threshold occurs both in males (in blue) and females (in pink). Interestingly, the depletion of the P2X4 gene in neurons increases the C-fiber threshold at levels similar to or even higher than those observed under physiological conditions, alleviating the hyperexcitability usually observed after neuropathic lesions. Mann-Whitney tests. **(C)** The intensity-response curve indicates how peripheral afferents respond to different intensities, reflecting the level of peripheral sensitization (sensitization of C-fiber activation of WDR neurons). In neuronal P2X4KO mice, under a neuropathic condition, the number of C-fiber responses is significantly lower than in WT mice at the same stimulation intensity, both in males and females. Mixed-effects analysis with Geisser-Greenhouse correction, followed by Šídák’s multiple comparisons test.

**Table 6:** Effect of deletion of neuronal P2X4 on dorsal horn of the spinal cord neurons excitability in male and female mice 60 days after sham or SNI surgery. Values are expressed as mean ± SEM. n = 9-58 neurons recorded per group (N= 5-21 mice per group). C-fiber threshold, minimal current intensity to trigger a C-fiber response; #C-response S1/S2/S3/S4/S5, number of C-fiber responses to electrical stimulation from 1 to 5 mA; SNI, spared nerve injury; WT, wild type; SynP2X4KO, neuronal P2X4 knockout mice.

| **In vivo recordings** | **Male** |  |  |  | **Female** |  |  |  |
|---|---|---|---|---|---|---|---|---|
|  | **Sham** |  | **SNI** |  | **Sham** |  | **SNI** |  |
|  | **WT** | **SynP2X4KO** | **WT** | **SynP2X4KO** | **WT** | **SynP2X4KO** | **WT** | **SynP2X4KO** |
| C-fiber threshold | 0.97 ± 0.07 | 1.16 ± 0.08 | 0.68 ± 0.06 | 1.35 ± 0.08 | 1.19 ± 0.07 | 0.88 ± 0.09 | 0.89 ± 0.12 | 1.6 ± 0.12 |
| #C-response S1 | 1.96 ± 0.55 | 2.38 ± 0.67 | 3.53 ± 1.01 | 1 ± 0.35 | 1.69 ± 0.33 | 2.43 ± 0.69 | 3 ± 0.84 | 0.45 ± 0.26 |
| #C-response S2 | 3.14 ± 0.69 | 5.53 ± 0.92 | 5.92 ± 1.10 | 3.12 ± 0.66 | 3.13 ± 0.54 | 4.39 ± 0.64 | 7.2 ± 1.34 | 3.4 ± 1.16 |
| #C-response S3 | 4.43 ± 0.83 | 7.22 ± 1.32 | 6.56 ± 0.88 | 3.97 ± 0.74 | 3.77 ± 0.61 | 5.78 ± 1.06 | 9.22 ± 1.59 | 4.37 ± 1.10 |
| #C-response S4 | 4.94 ± 0.90 | 9.26 ± 1.86 | 7.96 ± 0.99 | 5.14 ± 0.85 | 3.94 ± 0.66 | 6.5 ± 0.91 | 10.7 ± 2.27 | 5.13 ± 1.05 |
| #C-response S5 | 5.44 ± 0.98 | 8.30 ± 1.49 | 9.20 ± 1.00 | 6.02 ± 0.96 | 4.94± 0.77 | 6.5 ± 0.98 | 9.77 ± 1.45 | 6.45 ± 1.64 |

We next assessed intensity-response relationship (IR) using supraliminar stimulations. In sham males, evoked responses were slightly, but not significantly, higher in synP2X4KO than in WT mice. Following SNI, however, responses were significantly lowered in SynP2X4KO compared to WT mice (**Fig. 5C**, left panels). A similar pattern was observed in females, with significantly reduced responses in SynP2X4KO SNI compared with WT SNI mice (**Fig. 5C**, right curves).

Together, these electrophysiological findings indicate that neuronal P2X4 deletion primarily reduces spinal WDR neuronal excitability following SNI. This reduction may contribute to the partial attenuation of persistent mechanical hypersensitivity observed in SynP2X4KO mice.

## Discussion

Chronic pain remains a major clinical challenge, in part because the mechanisms underlying its initiation and maintenance are incompletely understood. Among the molecular mechanisms implicated in neuropathic pain, P2X4 has emerged as a key player, particularly in spinal microglia ^6^. However, its expression over time and the contributions of distinct cellular populations to established pain remain poorly defined. In the present study, we combined immunohistochemical, behavioral, and electrophysiological approaches in the SNI model to investigate these questions, with particular emphasis on pain persistence and sex-dependent sensory and affective manifestations.

### Transient spinal microglial P2X4 upregulation contrasts with persistent pain hypersensitivity

Our immunohistochemical analysis revealed robust P2X4 upregulation in spinal microglia at D15 after SNI, consistent with previous studies implicating microglial P2X4 in the early phase of neuropathic pain ^17^. However, this increase was transient, with P2X4 immunoreactivity returning to baseline by D30 and remaining at this level at D60. In contrast, mechanical allodynia and weight-bearing deficits–persisted at both time points. This temporal dissociation suggests that persistent hypersensitivity does not require sustained, detectable P2X4 upregulation in spinal microglial and raises questions about the cellular and anatomical substrates of its contribution to pain maintenance.

### Microglial P2X4 contributes to both the development and maintenance of neuropathic pain

To distinguish the contribution of microglial P2X4 to pain development and maintenance, we used Cx3cr1^CreERT2^ : P2X4^flox/flox^ mice and administered tamoxifen at two distinct time points: 21 days before SNI or 40 days after injury, when hypersensitivity was already established. P2X4 deletion before SNI prevented mechanical allodynia and weight-bearing deficits at D30 and D60, supporting a critical contribution to pain development, consistent with previous findings ^17^ ^10^. In contrast, deletion initiated at D40 reversed established mechanical hypersensitivity and weight-bearing deficits by D60, indicating that the contribution of microglial P2X4 extends beyond the early phase after injury, in accordance with previous results showing that spinal microglia are essential for chronic pain ^26^.

This finding is particularly intriguing because spinal P2X4 immunoreactivity was no longer detectable in spinal microglia at D30 and D60. The effect of delayed microglial P2X4 deletion is therefore difficult to reconcile with a mechanism relying exclusively on sustained P2X4 upregulation in spinal microglia. A functionally relevant level of spinal microglial P2X4 below the detection threshold of immunohistochemistry cannot be excluded. Alternatively, microglial P2X4 may contribute to pain maintenance outside the spinal cord, potentially in supraspinal regions involved in pain modulation. Further studies examining its spatial distribution and function at chronic time points will be required to identify the anatomical substrates of this late contribution.

The effects of microglial P2X4 deletion were observed in both males and females, supporting a contribution to persistent sensory hypersensitivity in both sexes. These findings contrast with previous reports of a predominantly male-specific contribution of microglial signaling to neuropathic pain ^27^, suggesting that the involvement of microglial pathways may depend on the molecular mechanisms and temporal stages investigated.

### Neuronal P2X4 plays a modulatory role in neuropathic pain

Given the transient expression of P2X4 in spinal microglia, we next investigated whether neuronal P2X4 contributes to persistent mechanical hypersensitivity. Neuronal P2X4 deletion did not affect basal mechanical withdrawal thresholds, suggesting that neuronal P2X4 is not necessary for baseline mechanical sensitivity under the conditions tested. Following SNI, however, SynP2X4KO mice exhibited significant but partial attenuation of mechanical allodynia in both sexes.

Electrophysiological recordings at D60 provided a functional correlate for this behavioral effect. Neuronal P2X4 deletion increased the stimulation threshold required to evoke C-fiber-mediated responses and reduced suprathreshold responses in dorsal horn WDR neurons following SNI. These findings support a contribution of neuronal P2X4 directly to persistent spinal hyperexcitability, consistent with previous evidence for its functional involvement in nociceptive pathways ^11^ ^13^. Because P2X4 deletion was not restricted to spinal neurons, however, these effects cannot be attributed exclusively to receptors expressed within the dorsal horn. Moreover, constitutive neuronal deletion does not distinguish a contribution to the development of sensitization from a requirement for its maintenance.

Together, these results reveal distinct cellular contributions of P2X4 during persistent neuropathic pain. Neuronal P2X4 deletion was associated with reduced spinal neuronal hyperexcitability, whereas delayed microglial P2X4 deletion reversed hypersensitivity despite the absence of detectable spinal microglial P2X4 upregulation. The latter observation may reflect a contribution from microglia outside the spinal cord or from spinal expression below our detection threshold.

### Sex-dependent timing of affective disturbances accompanies persistent sensory hypersensitivity in both sexes

A notable finding of our study is the difference in the timing of sensory and affective manifestations between sexes. Anxiety- and depressive-like behaviors were detectable at D30 in WT males but only at D60 in WT females. Their progressive emergence is consistent with previous work by Yalcin et al., who reported the development of mood-related alterations over time after peripheral nerve injury, with anxiety-related behaviors preceding depressive-like behaviors ^2^. By D60, both sexes in our study exhibited similar levels of affective disturbances. This pattern contrasts with mechanical allodynia and weight-bearing deficits, which were already present in both sexes at D30.

Together, these findings suggest that sensory and affective consequences of neuropathic pain follow distinct temporal dynamics and may be differentially regulated between sexes. While P2X4-dependent mechanisms involved in sensory hypersensitivity appear to operate similarly in males and females, the mechanisms underlying the development of affective disturbances may be subject to sex-specific temporal dynamics.

Global P2X4 deletion also prevented the emergence of anxiety- and depressive-like behaviors in both sexes. However, because these mice were also protected from persistent pain hypersensitivity, our data do not establish whether P2X4 contributes directly to affective disturbances or whether their absence is secondary to reduced pain. Dissecting these possibilities will require approaches that distinguish sensory from affective effects and determine the respective contributions of neuronal and microglial P2X4.

### Distinct cellular contributions of P2X4 to persistent neuropathic pain

Taken together, our findings support distinct contributions of microglial and neuronal P2X4 to persistent neuropathic pain. Microglial deletion before SNI prevented long-term hypersensitivity, whereas deletion initiated after pain establishment reversed it by D60. These findings support a role beyond pain initiation, although they do not establish a continuous requirement throughout the intervening period. In contrast, neuronal P2X4 deletion partially attenuated mechanical allodynia and reduced WDR neuronal hyperexcitability, supporting a modulatory contribution to persistent sensitization. The different effects of these deletions do not establish a hierarchical relationship between microglial and neuronal P2X4. Whether neuronal P2X4 acts downstream of microglial signaling or contributes through parallel mechanisms remains to be determined. The complete protection from both sensory and affective disturbances in P2X4 KO mice underscores the central role of this receptor in chronic pain. However, the fact that spinal microglial P2X4 expression is transient, while its functional requirement persists, highlights the complexity of P2X4-mediated signaling and the need for further studies to identify the long-lasting downstream effectors that sustain chronic pain. The reversal of established mechanical hypersensitivity following delayed P2X4 deletion, together with previous evidence of analgesic effects of selective P2X4 antagonists in experimental pain models ^28^, further supports the therapeutic potential of targeting P2X4 in established pain.

## Conclusion

Our study identifies distinct microglial and neuronal P2X4 contributions to persistent neuropathic pain in male and female mice. Microglial P2X4 is essential for both initiation and maintenance, whereas neuronal P2X4 plays a modulatory role. Despite the transient expression of microglial P2X4 in the spinal cord, its functional requirement persists, and its delayed global deletion reverses established pain. Anxiety and depressive-like behaviors develop earlier in males than in females, but P2X4 is required for their emergence in both sexes. These findings provide a comprehensive framework for understanding the temporal, cellular and sex-specific contributions of P2X4 to chronic pain and its comorbidities, with important implications for the development of targeted therapies for chronic pain.

## Acknowledgements

We thank the following Montpellier Biocampus core facilities: Réseau des Animaleries de Montpellier (RAM) (Iexplore facility), and the imaging facility MRI.

## Author contributions

D.G, B.S, N.L and P.N performed the experiments and analyzed the data. P.F, E.BG and L.U conceived the study and designed the experiments. L.U. wrote the first version of the manuscript. All authors commented on the manuscript.

## Availability of data and materials

All data generated or analyzed during this study are included in this published article.

## Declarations

Ethics approval and consent to participate. All experiments followed European Union (Council directive 2010/63/UE) and institutional guidelines for the care and use of laboratory animals. The animal experiment protocols used in this study were approved by the ethical committee for animal experimentation of Languedoc Roussillon (CEEA-LR; APAFiS #22741), of Bordeaux (CEEA50, APAFIS, #50393) and the French Ministry of Research.

## Consent for publication

Not applicable.

## References

1. Costigan, M., Scholz, J. & Woolf, C. J. Neuropathic Pain: A Maladaptive Response of the Nervous System to Damage. Annu. Rev. Neurosci. 32, 1–32 (2009).

2. Yalcin, I. et al. A Time-Dependent History of Mood Disorders in a Murine Model of Neuropathic Pain. Biol. Psychiatry 70, 946–953 (2011).

3. Won, S., Park, K., Lim, H. & Lee, S. J. Sexual dimorphism in cognitive disorders in a murine model of neuropathic pain. Behavioral and Brain Functions 16, 1 (2020).

4. Inoue, K., Tsuda, M. & Koizumi, S. ATP- and adenosine-mediated signaling in the central nervous system: chronic pain and microglia: involvement of the ATP receptor P2X4. J Pharmacol Sci 94, 112–114 (2004).

5. Coull, J. A. et al. BDNF from microglia causes the shift in neuronal anion gradient underlying neuropathic pain. Nature 438, 1017–1021 (2005).

6. Ulmann, L. et al. Up-regulation of P2X4 receptors in spinal microglia after peripheral nerve injury mediates BDNF release and neuropathic pain. J Neurosci 28, 11263–11268 (2008).

7. Beggs, S., Trang, T. & Salter, M. W. P2X4R + microglia drive neuropathic pain. Nature Neuroscience vol. 15 1068–1073 Preprint at 10.1038/nn.3155 (2012).

8. Inoue, K. & Tsuda, M. Microglia in neuropathic pain: cellular and molecular mechanisms and therapeutic potential. Nat. Rev. Neurosci. 19, 138–152 (2018).

9. Schwab, J. M., Guo, L. & Schluesener, H. J. Spinal cord injury induces early and persistent lesional P2X4 receptor expression. J. Neuroimmunol. 163, 185–189 (2005).

10. Kohno, K. & Tsuda, M. Role of microglia and P2X4 receptors in chronic pain. Pain Rep. 6, e864 (2021).

11. Lalisse, S. et al. Sensory neuronal P2RX4 receptors controls BDNF signaling in inflammatory pain. Sci. Rep. 8, (2018).

12. Duveau, A., Bertin, E. & Boué-Grabot, E. Implication of Neuronal Versus Microglial P2X4 Receptors in Central Nervous System Disorders. Neuroscience Bulletin vol. 36 1327–1343 Preprint at 10.1007/s12264-020-00570-y (2020).

13. Bernier, L., Ase, A. R. & Séguéla, P. P2X receptor channels in chronic pain pathways. Br. J. Pharmacol. 175, 2219–2230 (2018).

14. Bertin, E. et al. Increased surface P2X4 receptors by mutant SOD1 proteins contribute to ALS pathogenesis in SOD1-G93A mice. Cellular and Molecular Life Sciences 79, 431 (2022).

15. Bertin, E. et al. Increased surface P2X4 receptor regulates anxiety and memory in P2X4 internalization-defective knock-in mice. Mol. Psychiatry https://doi.org/10.1038/s41380-019-0641-8 (2020) doi:10.1038/s41380-019-0641-8.

16. Sim, J. A. et al. Altered hippocampal synaptic potentiation in P2X4 knock-out mice. J Neurosci 26, 9006–9009 (2006).

17. Gilabert, D. et al. Microglial P2X4 receptors are essential for spinal neurons hyperexcitability and tactile allodynia in male and female neuropathic mice. iScience 26, 108110 (2023).

18. Zhu, Y. et al. Ablation of NF1 function in neurons induces abnormal development of cerebral cortex and reactive gliosis in the brain. Genes Dev. 15, 859–876 (2001).

19. Yona, S. et al. Fate Mapping Reveals Origins and Dynamics of Monocytes and Tissue Macrophages under Homeostasis. Immunity 38, 79–91 (2013).

20. Ulmann, L. et al. Up-regulation of P2X4 receptors in spinal microglia after peripheral nerve injury mediates BDNF release and neuropathic pain. Journal of Neuroscience 28, 11263–11268 (2008).

21. Jahn, H. M. et al. Refined protocols of tamoxifen injection for inducible DNA recombination in mouse astroglia. Sci. Rep. 8, 5913 (2018).

22. Parkhurst, C. N. et al. Microglia Promote Learning-Dependent Synapse Formation through Brain-Derived Neurotrophic Factor. Cell 155, 1596–1609 (2013).

23. Bergmann, P. et al. Generation and Characterization of Specific Monoclonal Antibodies and Nanobodies Directed Against the ATP-Gated Channel P2X4. Front. Cell. Neurosci. 13, (2019).

24. Meseguer-Beltrán, M., Sánchez-Sarasúa, S., Landry, M., Kerekes, N. & Sánchez-Pérez, A. M. Targeting Neuroinflammation with Abscisic Acid Reduces Pain Sensitivity in Females and Hyperactivity in Males of an ADHD Mice Model. Cells 12, 465 (2023).

25. Vieira, W. F. et al. Neuropathic pain, mood, and stress-related disorders: A literature review of comorbidity and co-pathogenesis. Neurosci. Biobehav. Rev. 161, 105673 (2024).

26. Echeverry, S. et al. Spinal microglia are required for long-term maintenance of neuropathic pain. Pain 158, 1792–1801 (2017).

27. Sorge, R. E. et al. Different immune cells mediate mechanical pain hypersensitivity in male and female mice. Nat. Neurosci. 18, 1081–1083 (2015).

28. Inoue, K. Nociceptive signaling of P2X receptors in chronic pain states. Purinergic Signal. 17, 41–47 (2021).

